# RummaGEO: Automatic Mining of Human and Mouse Gene Sets from GEO

**DOI:** 10.1101/2024.04.09.588712

**Authors:** Giacomo B. Marino, Daniel J. B. Clarke, Eden Z. Deng, Avi Ma’ayan

## Abstract

The Gene Expression Omnibus (GEO) is a major open biomedical research repository for transcriptomics and other omics datasets. It currently contains millions of gene expression samples from tens of thousands of studies collected by many biomedical research laboratories from around the world. While users of the GEO repository can search the metadata describing studies for locating relevant datasets, there are currently no methods or resources that facilitate global search of GEO at the data level. To address this shortcoming, we developed RummaGEO, a webserver application that enables gene expression signature search of a large collection of human and mouse RNA-seq studies deposited into GEO. To develop the search engine, we performed offline automatic identification of sample conditions from the uniformly aligned GEO studies available from ARCHS4. We then computed differential expression signatures to extract gene sets from these studies. In total, RummaGEO currently contains 135,264 human and 158,062 mouse gene sets extracted from 23,395 GEO studies. Next, we analyzed the contents of the RummaGEO database to identify statistical patterns and perform various global analyses. The contents of the RummaGEO database are provided as a web-server search engine with signature search, PubMed search, and metadata search functionalities. Overall, RummaGEO provides an unprecedented resource for the biomedical research community enabling hypothesis generation for many future studies. The RummaGEO search engine is available from: https://rummageo.com/.

## Introduction

The Gene Expression Omnibus (GEO) contains tens of thousands transcriptomics studies and over two million genome-wide gene expression samples collected by RNA-seq^1^. Such massive transcriptomics profiling corpus covers many organisms, disease conditions, drug treatments, genetic perturbations such as knockouts, knockdowns, and overexpression of genes across tissues, cell types, and cell lines. This transcriptomics data in GEO, however, can be difficult to search and reuse because it is mostly provided in raw FASTQ files format, and metadata about the conditions of each study, and samples within each study, have inconsistent formatting and follow different naming conventions^2^. Multiple attempts have been made to make GEO studies better searchable by standardizing and restructuring the GEO metadata. For instance, GEOmetadb provides an R package and accompanying SQLite database to query GEO datasets locally, improving the speed of querying and the accessibility of the GEO metadata^3^. Similarly, ReGEO utilized natural language processing (NLP) techniques to extract time-points and disease terms from GEO metadata, enabling users to search by these attributes, as well as many other attributes, embedded in the original metadata^4^. Another notable effort is MetaSRA^5^. MetaSRA maps GEO metadata about samples and studies to ontologies and dictionaries. These mappings facilitate a metadata search engine that is more useful for identifying samples and studies from across different studies in GEO.

Although these efforts make GEO metadata more accessible and searchable, these resources do not enable users to search the GEO database at the data level, nor they provide direct access to uniformly aligned samples and signatures from the processed GEO studies. Several efforts aim to uniformly align GEO RNA-seq transcriptomics samples and make these accessible to users for reuse. For example, Recount, in its third iteration called Recount3, uniformly aligned more than 750,000 human and mouse RNA-seq samples from GEO^1^, GTEx^6^, and The Cancer Genome Atlas (TCGA)^7^, enabling users to more easily investigate and compare gene expression profiles across these resources^8^. Recount3 processed datasets are served via an R shiny web-based data explorer and an R package. The GEO RNA-seq Experiments Interactive Navigator (GREIN), also uniformly aligned over 600,000 GEO RNA-seq samples from human, mouse and rat. GREIN processed data is served via an interactive R shiny app to investigate these studies and request new GEO studies to be aligned and added to the database^9^. We have developed the All RNA-seq and ChIP-seq sample and signature search (ARCHS4) resource^10^. ARCHS4 provides access to over 2 million human and mouse uniformly aligned RNA-seq samples from GEO. Another similar effort called Digital Expression Explorer 2 (DEE2) provides uniformly aligned RNA-seq data for human and mouse as well as other species^11^. All together, these projects provide valuable uniformly processed data from GEO and other resources, making such data more accessible and reusable. However, none of these resources provide the uniformly aligned data for search at the data or signature levels.

Differential gene expression signatures associated with these studies, however, must still be manually computed. This means users need to manually parse the metadata associated with each sample to determine proper groupings of samples, which can be a time consuming process when attempted for thousands of studies. Various efforts have attempted to automatically or manually compute signatures from GEO studies. The CRowd Extracted Expression of Differential Signatures (CREEDS) resource, for instance, provides manually curated and automatically generated gene, drug, and disease perturbation signatures extracted from GEO studies^12^. These signatures were created via a crowdsourcing project that provided participants access to the tool GEO2Enrichr^13^. GEO2Enrichr enables users to extract differentially expressed genes from GEO studies using a browser extension. After identifying the control and perturbation samples, users can submit the computed signatures to Enrichr for reanalysis. A related project, GEN3VA^14^, saves signatures extracted with GEO2Enrichr, and then makes these signatures available to the public as collections based on hashtags. The main limitation of GEO2Enrichr and GEN3VA is that they only work for processing data from microarray studies. There are many other projects that enable users to manually annotate GEO studies and their metadata in a user-friendly interface. For example, GEOMetaCuration provides users with an intuitive GUI to label and submit relevant metadata and keywords associated with GEO studies^15^. BioJupies^16^ enables users to select samples from GEO studies that were uniformly aligned by ARCHS4 to label control and perturbation conditions, compute differential expression, and perform a variety of analyses and visualizations using automatically executed Jupyter Notebooks in the cloud. In a similar vein, TidyGEO^17^ and iLINCS^18^ allow users to select GEO series, examine and label their metadata, as well as perform multiple data cleaning and filtering tasks followed by standard downstream analyses such as differential expression and pathway analysis. These bioinformatics web apps, while useful, still require users to manually search and select their studies and conditions. To enable more efficient, automatic, and large-scale analyses, automated label extraction (ALE) has attempted to label GEO metadata based on gene expression signatures using logistic regression, predicting tissue, age, and gender of samples that do not have such annotations^19^. Other projects, for example, Restructured GEO (ReGEO) have attempted to extract disease and timepoint information of samples using natural language processing (NLP)^4^. While numerous projects aim to facilitate standardization, manipulation, and exploration of GEO studies and samples, no resource currently exists to enable searching GEO at the data level.

However, in the past, several efforts have been established to serve GEO data in a more digestible format. For example, ExpressionBlast used regular expression to identify groups of samples and normalized expression across studies for microarray data^20^. Search-Based Exploration of Expression Compendium (SEEK) was developed to provide search for gene or gene sets across a subset of human microarray and RNA-seq studies uniformly processed from GEO^21^. Unfortunately, both ExpressBlast and SEEK are no longer available or have not been updated since 2015, respectively. A more recent effort called GENe Expression Variance Analysis (GENEVA) leveraged the data from ARCHS4 to semi-automatically identify and serve groups of samples from human studies^22^. The GENEVA website that hosted the data and the search engine to serve these processed datasets is also no longer publicly available. To facilitate this type of search, here we performed automatic identification and grouping of conditions of GEO samples from thousands of GEO studies, and then performed differential expression analysis producing over hundred thousand human and mouse signatures that are made available for search via a user-friendly web interface.

## Methods

### Identifying conditions and computing signatures

All the human and mouse GEO studies aligned by ARCHS4 with at least three samples per condition, and at least six samples in total for a specific study, were considered for inclusion into the RummaGEO database. Studies with more than 50 samples were discarded because such studies typically contain large patient cohort data that is not amenable for simple signature computation that compares two or more conditions. Samples were grouped using the metadata provided by each study. Specifically, K-means clustering of the embedding of concatenated sample_title, characteristic_ch1, and source_ch1 fields were used to classify the conditions. To create condition titles, common words across all samples for each condition were retained. The limma-voom R package^23^ was used to compute differential expression signatures for each condition against all other conditions within each study. Additionally, we attempted to first identify any control conditions based on metadata associated with each sample. To achieve this, a set of keywords that describe control conditions was compiled. The set of such terms contain, for example, “wildtype”, “ctrl”, and “DSMO”. If such terms were identified, they were used to compare the samples labeled with such terms to all other condition groups. Up and down gene sets were extracted from each signature for genes with an adjusted p-value of less than 0.05. If less than five genes met this threshold, the gene set was discarded. If more than 2000 genes met this threshold, the threshold was lowered incrementally from 0.05 to 0.01, then 0.005, and lastly 0.001, until less than 2000 genes were retained.

### Data level silhouette scores

For each study with identified conditions based on metadata clustering, a silhouette score was computed to determine the data-level adherence to the assigned groupings based on the metadata. All aligned counts were extracted from ARCHS4^10^ and normalized by the number of reads aligned, followed by log2 transformation, as well as quantile and z-score normalization. Principal component analysis (PCA) was then performed on the normalized data, and the silhouette scores were computed from the distance between the samples in each condition in the two-dimensional PCA space. Silhouette scores range from -1 to 1, where a value of 1 would indicate perfect clustering and -1 would indicate fully disjointed clusters.

### Search engine implementation

Given the large number of gene sets contained within the RummaGEO database, we employ a fast search engine strategy implementation previously described for Rummagene, a web-server application that hosts over 700,000 gene sets extracted from supporting materials of publications listed on PubMed Central^24^. Please see the methods section in this publication about the search engine strategy employed.

### Identifying functional terms from sample and study metadata

Functional terms were extracted from both sample (GSM) and study (GSE) metadata. These functional terms include: tissues, cell types, and cell lines (these terms were associated with the BRENDA Tissue Ontology^25^); diseases and phenotypes (these terms were sourced from DisGeNet^26^); drugs and small molecules (these terms were associated with International Chemical Identifiers (InChI) keys); and genes and proteins (these terms were associated with NCBI^27^ gene symbols). Synonyms and official terms were retained and associated with each study (GSE).

### Co-occurrence gene-gene similarity matrix

Gene-gene co-occurrence matrices were computed for coding genes (19,484 for human, and 22,350 for mouse) and non-coding genes (41,366 for human and 29,143 for mouse) using 50,000 randomly selected RummaGEO gene sets. The co-occurrence probabilities for any two genes *P*(*α, β*) were computed as previously described by Clark and Ma’ayan^28^. Using the co-occurrence matrix of human coding genes, we then computed the cosine similarity, Jaccard similarity, and normalized pointwise mutual information (NPWMI) between all pairs of genes as follows:

- 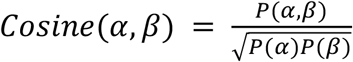
- 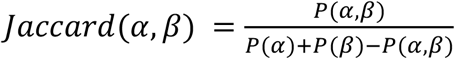
- 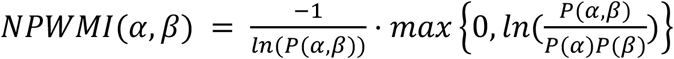

### Benchmarking transcription factor and kinase enrichment analyses

Identification of official NCBI gene symbols contained within GSE and GSM metadata enabled the creation of consensus transcription factor and kinase gene set libraries. Any signatures with a kinase or transcription factor term included in the GEO metadata were included. Each gene set for transcription factors and kinases, respectively, is formed by taking the intersection of genes identified in at least 5% of the gene sets mentioning the given transcription factor or kinase. The benchmarking datasets previously employed for the ChEA3^29^ and KEA3^30^ resources were used to assess the quality of the transcription factors and kinases RummaGEO libraries in regards to identifying the “correct” transcription factor and kinase. The Fisher’s exact test was applied to perform the enrichment analysis to rank transcription factors or kinases by p-value in each benchmarking dataset. When generating the ROC curves, the negative class was down-sampled to an equal size as the positive class. Downsampling was randomly performed over 5,000 iterations, and the mean ROC curves and AUCs were reported. To generate the composite ROC curves for each benchmarking library, the numpy *interp* function was utilized enabling linear interpolation of the generated points from the 5,000 ROC curves.

### Extracting key terms from abstracts

To enrich the metadata of RummaGEO gene sets with descriptive terms from biomedical text, the human and mouse GSEs included in RummaGEO were annotated with three different types of key terms based on the published paper associated with each GSE. The GEO DataSets database was first queried for the earliest PubMed ID linked to each GSE, and the corresponding article information was then retrieved from the PubMed database using the NCBI’s E-utilities Esearch function. In total, 13,427 human studies were annotated with data from 8,804 unique articles, and 15,478 mouse studies were annotated with data from 10,269 unique articles. For 53.5% of the human studies, and for the 54.0% of the mouse studies, the PubMed metadata included key terms provided by the authors and/or the publishing journals. Another source of key terms is the Medical Subject Headings (MeSH) thesaurus, a controlled vocabulary produced by the National Library of Medicine (NLM). MeSH headings were found for 87.7% of the human studies, and 92.0% of the mouse studies. A third set of key terms was generated for all studies in RummaGEO by submitting the abstracts from each article associated with each study to the Mistral-7B-Instruct-v0.2 LLM^31^, accessed via the HuggingFace API. The LLM was presented with the abstract text and prompted to return up to ten biomedical key terms. Without processing any terms, the human key term collection included 16,488 unique PubMed key terms; 7,935 unique MeSH headings; and 48,462 unique LLM-generated key terms. The mouse key terms comprised 17,742 unique PubMed key terms; 8,212 unique MeSH headings; and 52,152 unique LLM-generated key terms.

All terms were first normalized to a standard capitalization, punctuation, and grammatical forms. Due to the unstructured nature of the PubMed key terms and LLM-generated key terms, these two collections were further processed to consolidate key terms that are semantically synonymous or similar. To organize synonymous terms without introducing new terminology, a stemming-like procedure was implemented to identify small clusters of terms (<5) with high textual overlap. Each term in the cluster was replaced with the most frequent term, and these substitutions were assessed manually to ensure semantic equivalency. Terms that are overly general were then filtered out manually. The final processed collection of human key terms includes 10,713 unique PubMed key terms; 6,506 unique MeSH headings; and 31,210 unique LLM-generated key terms. The mouse processed key term collection contains 11,587 unique PubMed key terms; 7,140 unique MeSH headings; and 33,302 unique LLM-generated key terms.

### Term Categorization

To enable domain-specific enrichment analyses, RummaGEO key terms were sorted into four categories using Mistral-7B-Instruct-v0.2^31^. The categories were defined as follows: Category A: disease, phenotype; Category B: gene, metabolite, protein, drug, lipid, RNA, variant, receptor; Category C: organism, organ, tissue, cell line, cell type, organelle; Category D: pathway, biological process, family of genes, chromosome. The LLM was provided with a description of each of these categories and was asked to select the most appropriate choice for each key term. For cases where the LLM responses were uncertain, the term was categorized manually. Out of all human and mouse key terms, 5.4% were categorized as A, 33.3% as B, 19.5% as C, 32.5% as D, and 9.3% as ‘other’ (Table S2).

### Assigning enrichment terms to gene sets with Enrichr

To provide additional metadata and functional terms for RummaGEO gene sets, enrichment analysis is precomputed for each gene set for seven Enrichr^32^ libraries: ChEA 2022, KEGG 2021 Human, WikiPathway 2023 Human, GO Biological Process 2023, MGI Mammalian Phenotype Level 4 2021, Human Phenotype Ontology, and GWAS Catalog 2023. Significance is assessed using Fisher’s exact test and adjusted p-values are computed with Benjamini–Hochberg method^33^. Only significant terms with an adjusted p-value of <0.05 were retained. To assess the significance of the Enrichr terms’ appearance in the RummaGEO gene set search page, the Kolmogorov–Smirnov test is utilized, comparing the distribution of sets that are significantly enriched for that term compared to a uniform distribution. Enrichr terms are in the top 500 enriched sets returned from the search.

### Term enrichment analysis

Term enrichment is provided for the LLM extracted functional terms described above for four categories: Category A: disease, phenotype, Category B: gene, metabolite, protein, drug, lipid, RNA, variant, receptor; Category C: organism, organ, tissue, cell line, cell type, organelle; and Category D: pathway, biological process, family of genes, chromosome. Since functional terms are extracted per study (GSE), we compute term significance utilizing Fisher’s exact test on categorized terms from the first 5,000 unique GSEs returned from the RummaGEO gene set search and additionally adjusted p-values are computed with the Benjamini–Hochberg method.

### Global visualization of signatures with UMAP

To integrate the human and mouse gene sets from GEO, all genes were mapped to uppercase and only protein coding genes were retained. Gene sets were then converted to inverse document frequency (IDF) one-hot vectors for each set using the Scikit-learn package^34^. We then utilized truncated Singular Value Decomposition (SVD)^35^ to reduce the dimensionality of the IDF vectors to the largest 50 singular values. Then, to convert the vectors into two-dimensional space, UMAP^36^ was applied with the default parameters.

### Hypothesis generation with GPT

To generate hypotheses relevant to the user-inputted gene set, we utilize the OpenAI chat completion API. The user is required to submit a word description of their submitted gene set in the form of a summary or an abstract. RummaGEO takes this description together with the matching RummaGEO gene set study abstract, and the top three significantly enriched terms from the overlapping genes from the following Enrichr^37^ libraries: WikiPathway 2023 Human, GWAS Catalog 2023, GO Biological Process 2023, and MGI Mammalian Phenotype Level 4 2021. The prompt additionally instructs the large language model (LLM) to reference all the provided descriptions and contexts of the gene sets, as well as the highly enriched terms from Enrichr. Hypotheses are then parsed to find references for any of the enriched terms, and insert the enrichment statistics as part of the hypothesis description output.

## Results

### Descriptive statistics of the contents within the RummaGEO database

The initial release of RummaGEO contains 135,264 human and 158,062 mouse gene sets extracted and processed from 23,395 GEO studies. In general, most genes appear in only a small number of gene sets in both the collections of human-(Fig. 1A) and mouse-gene sets (Fig. 1D). There are many gene sets with less than 100 genes while the remaining sets are relatively equally distributed for both human (Fig. 1B) and mouse (Fig. 1E). The maximum gene set size that we defined is 2,000. Additionally, while the majority of the studies contributed just a few gene sets, there are periodic peaks for studies that contributed more sets. This periodicity is a result of the possible combinations of conditions and groups with a bias toward having an even number of groups in the study design (Figs. 1C-F). By identifying functional terms in sample metadata, we found that a large majority of studies (GSEs) contain the tissue or the cell type term which is often found in the *source_ch1* metadata field. Diseases also were identified for over half of the human GSEs (6,550), but much less disease terms were identified for mouse GSEs (1,091). Additionally, smaller subsets of GSEs also mentioned genes, drugs, and cell lines in the sample or the study metadata (Fig. 1G-H). From the identified gene symbols, we also extract the subsets of kinases and transcription factors (Figs. 1I-J). When plotting the contribution of studies over time, we observe a linear increase in GSEs added to RummaGEO. The drop in GSEs that started in 2022 due to processing data from the 2023 release of ARCHS4 (Fig. 1K). Silhouette scores were computed on the dimensionality reduced samples to examine whether the samples cluster in expression space as expected based on the groupings determined by the metadata. The silhouette scores for both human and mouse exhibit a bimodal distribution with one peak where the metadata and data levels are highly aligned (~0.5) and another peak where there is less alignment between the data and metadata (~-0.1) (Fig. 1L).

**Fig. 1.**
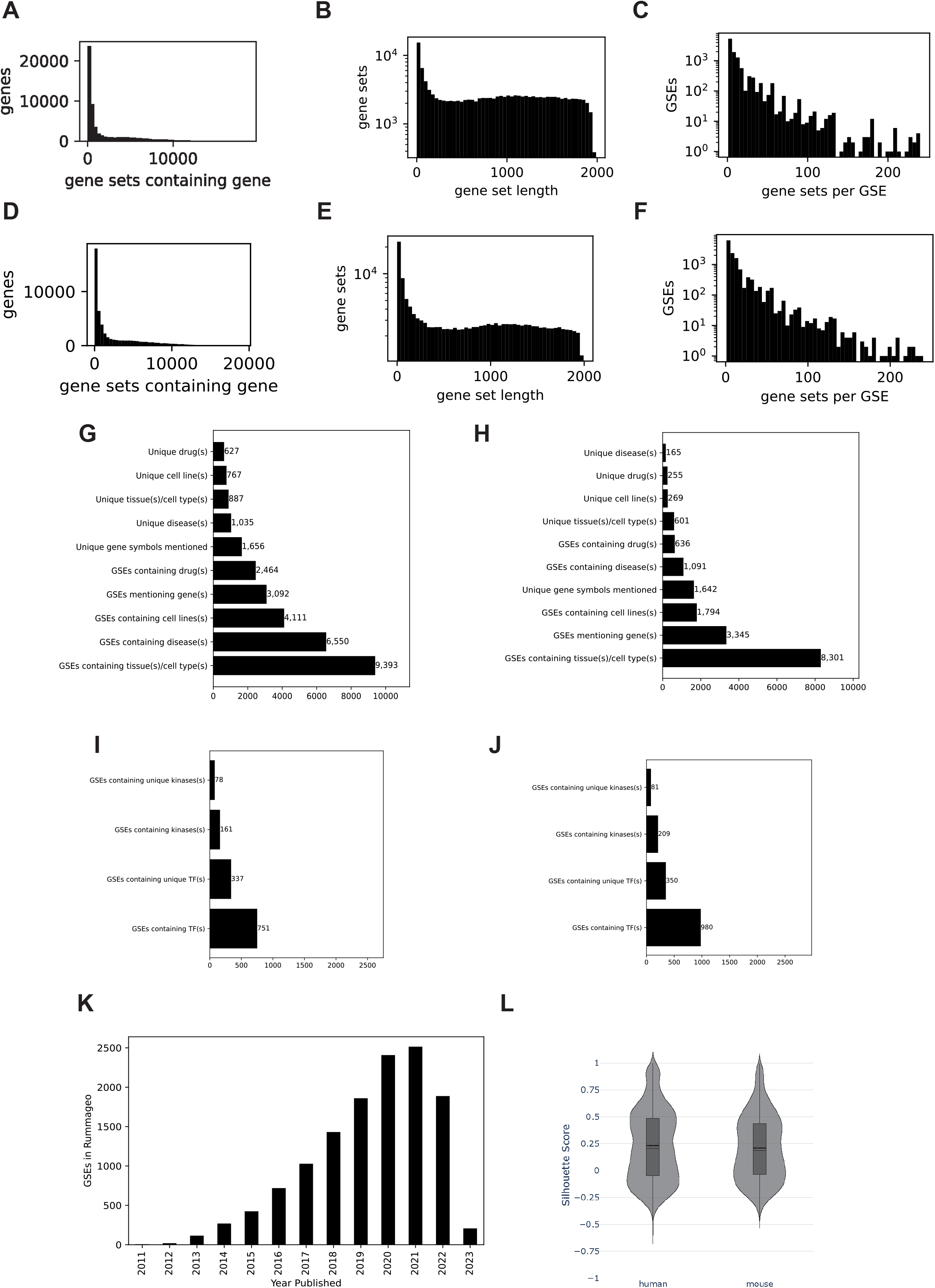
Distributions of genes and gene sets within the RummaGEO database. **A**. Distribution of genes across the human gene sets; **B**. Distribution of gene set lengths across the human gene sets; **C**. Human gene sets per GSE; **D**. Distribution of genes across mouse gene sets; **E**. Distribution of gene set lengths across mouse gene sets; **F**. Mouse gene sets per GSE; **G**. GSEs per year; **H**. Silhouette scores across all human and mouse studies; **I**. Unique and non-unique diseases, drugs, cell lines, tissues, and cell types, and genes mentioned in GSE and GSM metadata for human studies. **J**. Unique and non-unique diseases, drugs, cell lines, tissues and cell types, and genes mentioned in GSE and GSM metadata for mouse studies. **K-L**. Unique and non-unique kinase and transcription factors identified from gene mentions in GSE and GSM metadata for human (**K**) and mouse (**L**) studies.

### Global visualization of gene sets with UMAP

To visualize all of the gene sets within the RummaGEO database, gene sets were vectorized and visualized as a UMAP. Despite the harmonization of human and mouse gene symbols, we observe significant separation of the human and mouse gene sets in the global UMAP. On the other hand, the up and down signatures within each species are highly mixed (Fig. 2A). Next, we used the metadata extracted from the GSEs and GSMs to color the gene sets in the UMAP with the aim of elucidating additional patterns. When coloring by the most common tissues, we observe coherent groups of samples (Fig. 2B), indicating that the gene sets in RummaGEO are significantly influenced by their tissue of origin. Although less pronounced, the gene sets also appear to group by disease when visualizing the top 10 identified diseases (Fig. 2C). This is particularly apparent for diseases such as leukemia and lymphoma possibly due to their blood and bone marrow origins.

**Fig. 2.**
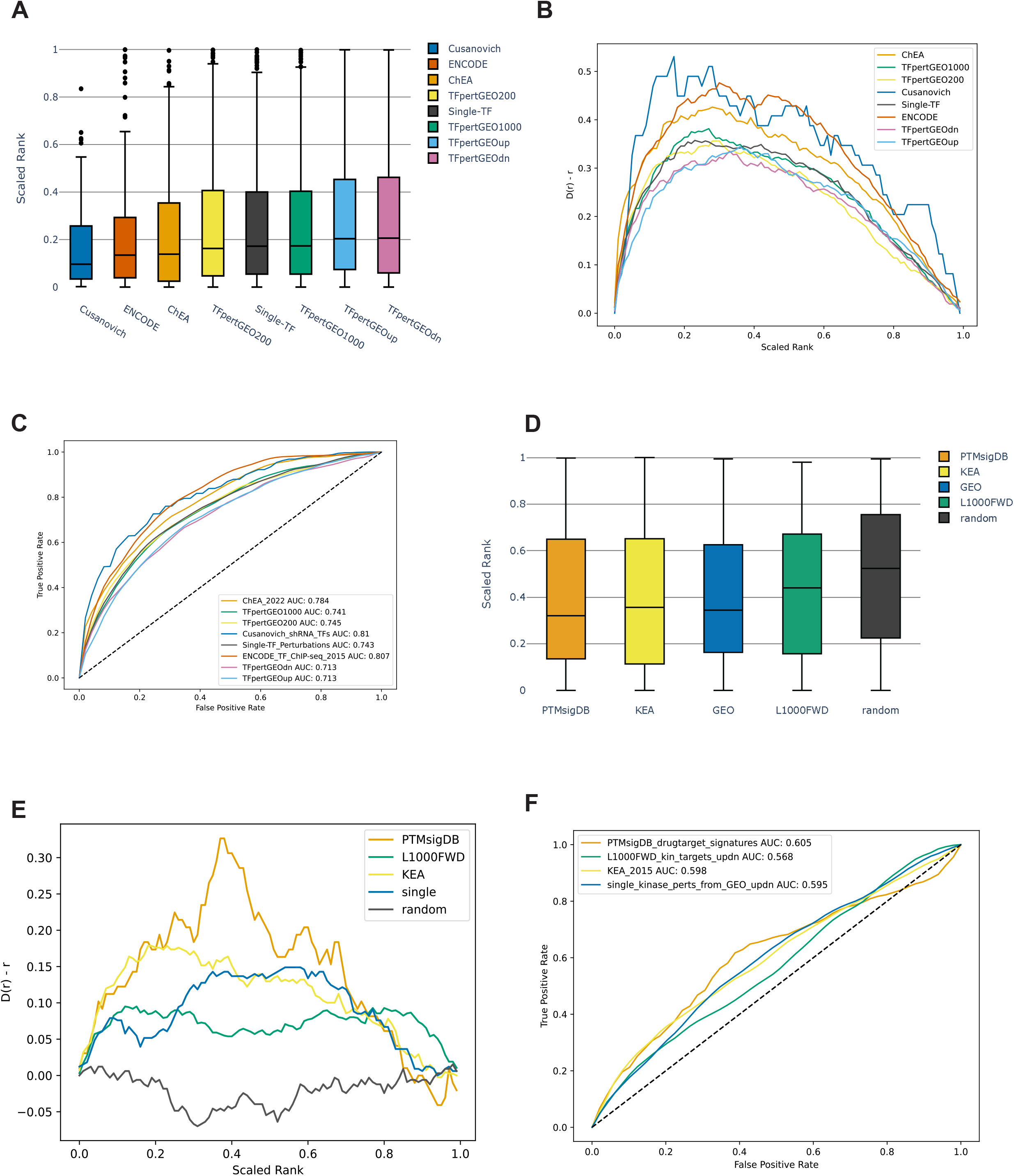
UMAP projection of all human and mouse gene sets in the RummaGEO database. **A**. Human up, human down, mouse up, and mouse down gene sets colored separately; **B**. The same UMAP is colored by the top 10 most mentioned tissues in the GSEs and GSMs metadata; **C**. The UMAP is colored by the top 10 most mentioned diseases in GSEs and GSMs metadata.

### Comparison of the RummaGEO and Enrichr gene set spaces

To better understand the breadth of coverage of the RummaGEO gene sets, we compared it to the Enrichr^37^ gene set space. Enrichr has thousands of curated gene sets spanning several domains and categories including: transcription, pathways, ontologies, diseases and drugs, cell types and tissues, miscellaneous, and crowd generated. Enrichr gene sets mainly cluster by category (Fig. 3A). Overlaying the RummaGEO gene sets on top of the Enrichr gene sets, we observe that the majority of gene sets in RummaGEO fall on the ‘crowd’ category. This is expected because the crowd category is also mainly composed of user extracted gene sets from GEO studies (Fig. 3B). There is also some overlap between Enrichr and RummaGEO gene sets in the center of the UMAP. However, these sets belong to several Enrichr categories so it is difficult to discern a clear pattern from that region of the UMAP.

**Fig. 3.**
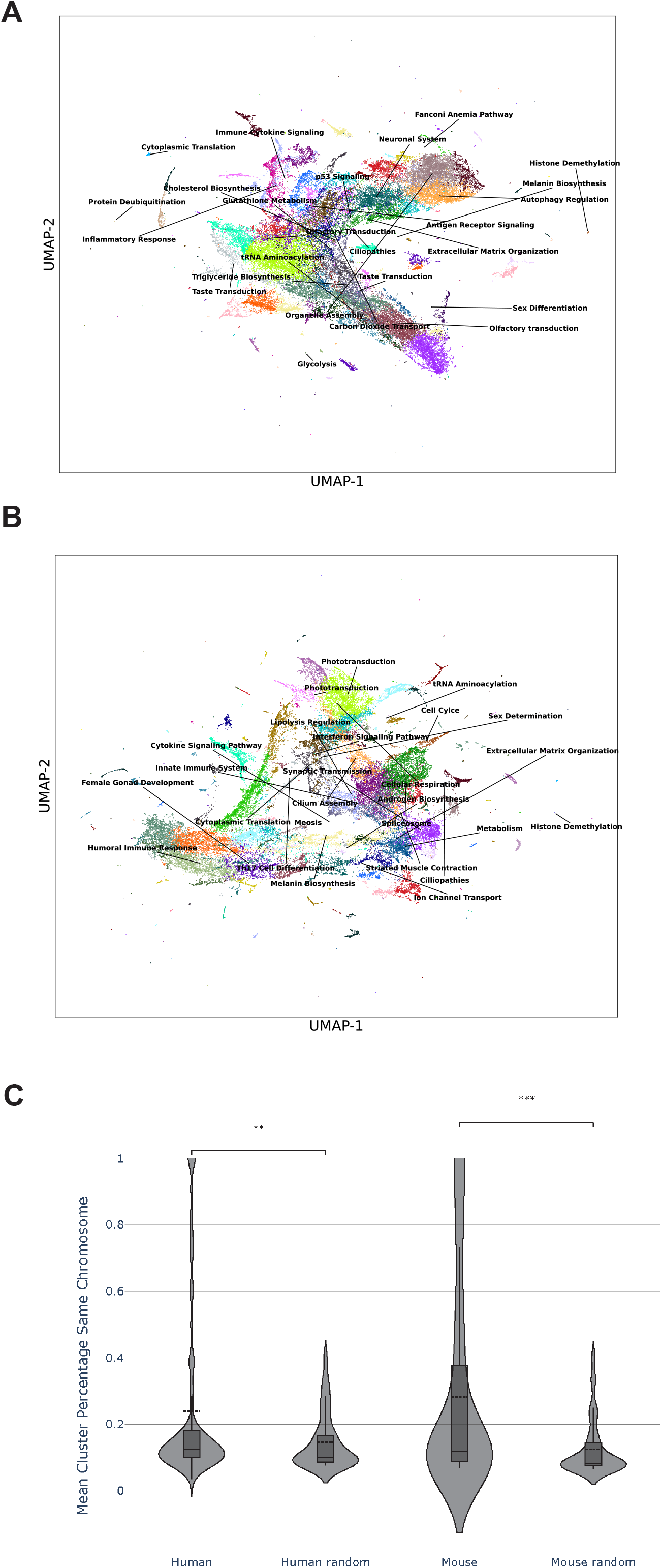
UMAP projection of the Enrichr and RummaGEO gene set spaces. **A**. Enrichr gene set space, colored by Enrichr categories; **B**. RummaGEO human and mouse gene sets mapped to human protein coding genes overlaid onto the Enrichr gene set space.

### Global visualization of genes with UMAP

In addition to examining the RummaGEO gene set space, we can also transpose the matrix to examine the gene similarity space. By plotting the gene vectors into two dimensions, and then automatically identifying clusters of genes, we can identify functional clusters, and see if groups of genes are differentially expressed by their chromosomal locations. First, we automatically identified clusters using the Leiden algorithm^38^. The algorithm identified 77 human, and 67 mouse clusters (Figs. 4A and 4B). For more than two-thirds of the mouse clusters, and for over half of human clusters, we identify significant functional terms with Enrichr^37^, these include GO Biological Processes^39^, KEGG^40^, Reactome^41^, WikiPathway^42^ (Table S1). This approach identified clear functional modules such as innate immune response, cytokine signaling, cell cycle, and regulation of autophagy. Additionally, to assess the influence of chromosome location on the gene clustering, we computed the percentage of genes originating from the same chromosome in each cluster (Fig. 4C). We observe that for both the human and mouse clusters, there are some groupings that exhibit enrichment for specific chromosomal locations.

**Fig. 4.**
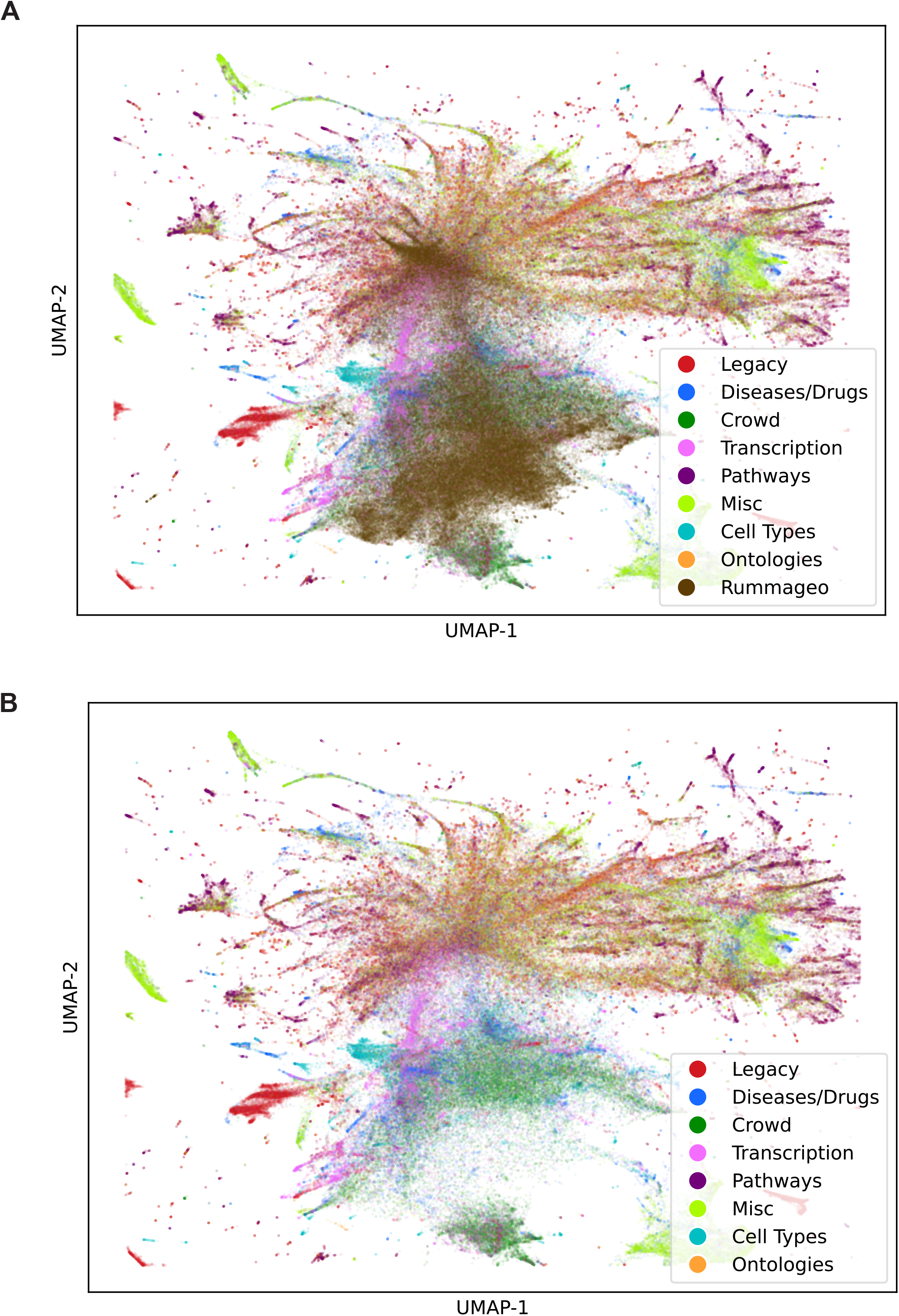
UMAP projection of all human and mouse genes in the RummaGEO database. **A**. UMAP projection of human genes clustered with the leiden algorithm; **B**. UMAP projection of mouse genes clustered with the Leiden algorithm; **C**. Fraction of genes from the same chromosome in human and mouse clusters and at random.

### Benchmarking transcription factor and kinase libraries created from the RummaGEO database

To assess the ability of RummaGEO signatures to recover known transcription factor targets and kinase substrates, we created transcription factor and kinase libraries from the RummaGEO gene-gene co-occurrence matrix for coding genes. In order to benchmark these libraries, we utilized ChEA3^29^ and KEA3^30^ and the benchmarking libraries these resources previously collected. Enrichment analysis was performed using the RummaGEO transcription factors and kinases libraries, and the rank of a given transcription factor or a kinase from the benchmarking datasets was determined by the significance of the overlap based on the p-value from the Fisher’s exact test. For transcription factors, the Cusanovich shRNA TFs^43^ library showed the most accurate recovery of transcription factors (area under the curve (AUC): 0.81) (Figs. 5A-5C). All other benchmarking libraries showed greater than 0.70 AUC. For the kinase libraries, PTMsigDB^44^ drug signatures showed the highest AUC of 0.605 (AUC: 0.598) (Fig. 5 D-F). The lower performance for kinases is expected because the RummaGEO gene-gene co-expression matrix is based on mRNA expression, while kinase phosphorylation events happen at the proteome and phosphoproteome levels. Signatures from transcription factor and kinase perturbations extracted from GEO did not show the highest recovery. This is less expected because the RummaGEO kinase and transcription factor libraries originate from the same source. Comparing the performance of the RummaGEO transcription factor and kinase libraries to those created from Rummagene^24^, the RummaGEO library performs similarly, with transcription factor recovery being slightly better, and kinase recovery slightly worse as expected.

**Fig. 5.**
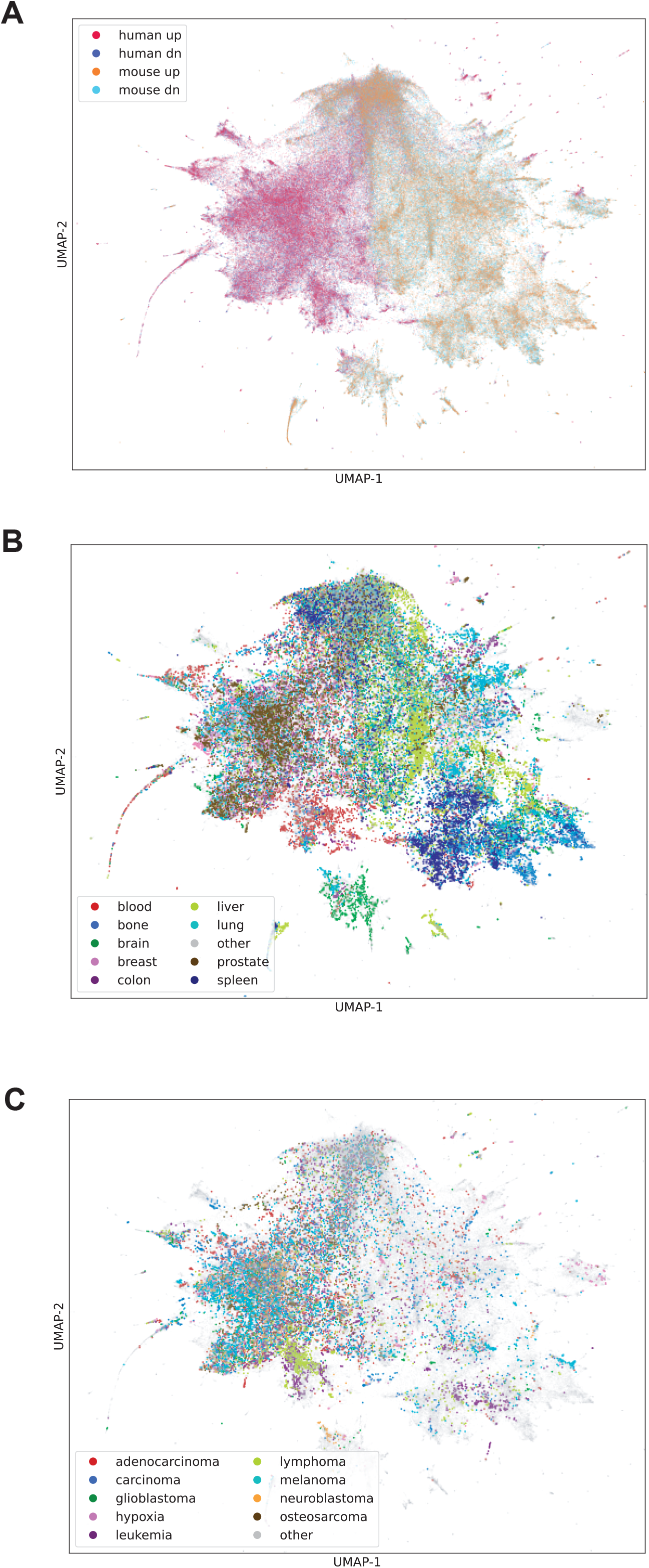
RummaGEO kinase and transcription factor libraries benchmarking. **A**. Scaled rank (0 highest rank, 1 lowest rank) for transcription factor benchmarking libraries as computed by the Fisher’s exact test; **B**. Deviation of the cumulative distribution for scaled ranks of each transcription factor from uniform distribution (Kolmogorov-Smirnov test for goodness of fit compared to uniform distribution: ChEA (ChEA_2022) P = 2.20E-88; TFpertGEO1000 P = 4.40E-115; TFpertGEO200 P = 6.84E-119; Cusanovich_shRNA_TFs P = 2.05E-12; Single-TF_Perturbations P = 2.98E-117; ENCODE_TF_ChIP-seq_2015 P = 1.14E-110; TFpertGEOdn P = 1.35E-89; TFpertGEOup P = 3.32E-92; **C**. 5,000 bootstrapped curves with downsampled negative class were generated to compute mean receiver operating characteristic (ROC) curves and mean area under the ROC curves (AUC) for transcription factors. **D**. Scaled rank (0 highest rank, 1 lowest rank) for kinase benchmarking libraries as computed by Fisher’s exact test; **E**. Deviation of the cumulative distribution for scaled ranks of each kinase from uniform distribution (Kolmogorov-Smirnov test for goodness of fit compared to uniform distribution: PTMsigDB_drugtarget_signatures P = 1.66E-06; L1000FWD_kin_targets_updn P = 1.09E-08; KEA_2015 P = 9.39E-10; single_kinase_perts_from_GEO_updn (single) P = 5.94E-08; **F**. 5,000 bootstrapped curves with downsampled negative class generated to compute mean receiver operating characteristic (ROC) curves and mean area under the ROC curves (AUC) for kinases.

### Generating hypotheses with GPT-4

After submitting a gene set for analysis with RummaGEO and examining the top matching results, the user can provide a textual description of the gene set they submitted, and then generate hypotheses with GPT-4, an LLM model developed by OpenAI. Once we obtain the descriptions of both the query gene set, the matching gene set from the RummaGEO database, as well as significantly enriched terms from their overlapping genes from Enrichr^37^, we can prompt an LLM to produce plausible reasoning for the observed highly significant set overlap. Although caution should be exercised in interpreting the hypotheses generated by the LLM, these hypotheses can provide valuable insight and assist with initial reasoning for explaining the observed overlap between the two sets. We outline several use cases below to demonstrate this feature of RummaGEO:

#### Use case 1: Investigating dysregulated mechanisms in Alzheimer’s disease (Rummagene PMC5534941-tp2017110x1.docx-6-Alzheimer_s_disease; n=42)

Searching “Alzheimer’s” in Rummagene^24^ provides numerous gene sets extracted from supporting materials of articles deposited in PMC related to this disease. One such gene set examines the genetic risk factors shared between coronary artery disease and Alzheimer’s disease (AD). In Supporting Table 3, the authors list a set of genes known to have association with AD based on published genetics studies.

https://rummagene.com/pubmed-search?page=1&q=PMC5534941-tp2017110x1.docx-6-Alzheimer_s_disease

Submitting this gene set to RummaGEO, results in the most significantly overlapping gene set to come from a study that inhibits STAT5 in acute myeloid leukemia (AML).

https://rummageo.com/enrich?dataset=eb388fc4-6ead-46d5-93c1-28c56548ee5d

To further investigate why these two gene sets might be related, we generated an hypothesis using the abstract of the PMC article from which the gene set was sourced as the description. From the descriptions of both gene sets and the significant Enrichr terms from their overlapping genes, GPT-4 produces a plausible explanation for the highly significant overlap:

*The high overlap between the user-submitted gene set and the GEO gene set could be due to the shared involvement of these genes in lipid metabolism and cholesterol transport, as well as their association with disease states such as acute myeloid leukemia (AML) and Alzheimer’s disease (AD)*.

The hypothesis describes both gene sets utilizing the provided descriptions:

> *The GEO gene set is derived from a study investigating the role of STAT5 in AML, particularly its activation by FLT3-ITD, a constitutively active tyrosine kinase. The study also explores the potential of a novel inhibitor, AC-4-130, in disrupting STAT5 activation and thereby impairing the proliferation and growth of AML cells*.
>
> *On the other hand, the user-submitted gene set is based on a study examining the genetic overlap between coronary artery disease (CAD) and AD, as well as the shared risk factors between these two diseases. The study found that genetic susceptibility to CAD modifies the association between cardiovascular disease (CVD) and dementia, likely through associations with shared risk factors*.

Then the hypothesis describes how the enrichment of their overlapping genes supports the shared mechanism related to lipid metabolism and cholesterol transport.

> *The enriched terms from the overlapping genes of the two sets further support this hypothesis. The terms Statin Inhibition Of Cholesterol Production WP430, Fatty Acids And Lipoproteins Transport In Hepatocytes WP5323, and Cholesterol Metabolism WP5304 from WikiPathway_2023_Human suggest a shared involvement in lipid metabolism and cholesterol transport. This is further supported by the GO_Biological_Process_2023 terms Phospholipid Efflux (GO:0033700), Cholesterol Efflux (GO:0033344), and Cholesterol Transport (GO:0030301)*.

Several studies in the literature, including the selected study, already support the hypothesis that there is accumulation of cholesterol^45^ and dysregulation of lipid metabolism^46^ in AD, and cholesterol metabolism reprogramming^47^ and lipid metabolism reprogramming^48^ is also happening in AML. The high overlap between the AD genes and genes from AML also further support the link between AD and inflammation. Thus, through this approach, we identify a dysregulated mechanism shared between AD and AML pointing to the key regulator STAT5. STAT5 activation has been reported to be protective in a mouse model of AD^49^.

#### Use case 2: Relation of Senescence Related Genes and Ewing’s Sarcoma Tumors

To identify GEO studies related to cellular senescence, a gene set was sourced from SenoRanger, which identified 301 genes highly expressed in senescent cells and lowly expressed in healthy human cells and tissues using a computational screen^50^.

https://rummageo.com/enrich?dataset=cee77176-0611-40b8-8a97-26c870d5c363

The most significantly overlapping signature identified by RummaGEO is related to the analysis of Ewing’s sarcoma family of tumors (ESFT) cell lines and the dysregulation of EWSR1 and BRCA1 genes^51^. The hypothesis theorizes that there are shared biological pathways between Ewing’s sarcoma and senescence, particularly those related to extracellular matrix organization, and the structure and strength of connective tissue:

> *The terms* “*Extracellular Matrix Organization (GO:0030198)"*, “*Extracellular Structure Organization (GO:0043062)", and* “*External Encapsulating Structure Organization (GO:0045229)” from GO_Biological_Process_2023 suggest that both senescent cells and ESFT cells may undergo changes in their extracellular matrix and structure, possibly as a response to stress or as a mechanism to evade immune surveillance*.
>
> *Finally, the terms* “*abnormal cutaneous collagen fibril morphology MP:0008438"*, “*decreased skin tensile strength MP:0003089", and* “*abnormal tendon morphology MP:0005503” from MGI_Mammalian_Phenotype_Level_4_2021 suggest that both senescence and ESFT may affect the structure and function of connective tissues, possibly due to alterations in extracellular matrix organization and structure*.

Senescence and its role and relation to extracellular matrix organization is actively being investigated^52,53^. Interestingly, the Ewing’s sarcoma gene, Ews, has been observed to be essential in modulating senescence in hematopoietic stem cells^54^. Given the relation of these sets, and the enriched terms related to their overlap, the hypothesis identifies ECM organization as the central theme of the apparent overlap. It should be noted that SenoRanger was created by extracting gene sets from several published studies that induced fibroblast cell lines to undergo senescence. Hence, the enrichment for ECM organization when combining Ewing’s disease signatures with fibroblasts undergoing senescence may provide actionable clues towards the identification of novel therapeutic strategies for Ewing’s sarcoma.

### The RummaGEO search engine

The RummaGEO website supports four main components of functionality and search. The first is gene set enrichment which utilizes an in-memory algorithm^24^ to calculate Fisher’s exact test results quickly. Enriched signatures can be filtered by a term and by a silhouette score threshold. For each significantly enriched signature, users may also generate an hypothesis with GPT-4 as described above. In addition to returning significantly overlapping gene sets, RummaGEO also provides term enrichment from three sources: MeSH terms^55^, PubMed key terms, and terms extracted by an LLM. Once a gene set has been enriched, the enriched terms are available as an additional tab, wherein users are provided with a bar chart, a table, and a wordcloud created from the enriched terms. The RummaGEO database can be queried in conjunction with PubMed to find signatures from GEO studies associated with any returned publications from a PubMed search. Users can additionally search RummaGEO through the GEO metadata to find signatures associated with any search term. Finally, the human and mouse gene set libraries and accompanying metadata are available to download from the RummaGEO website. Additionally, users can learn more about how to use RummaGEO and the available API from the ‘About’ and ‘Documentation’ pages.

## Discussion and Conclusions

By automatically identifying conditions from uniformly aligned studies from GEO and computing differential gene signatures, we were able to produce 135,264 human and 158,062 mouse gene sets extracted from 23,395 GEO studies. These sets provide differential expression knowledge across a wide array of experimental conditions. It should be noted that the identified gene sets come in pairs of up and down sets for each condition. Hence, if we term the paired up and down sets as signatures, there are 67,632 human and 79,031 mouse signatures in the RummaGEO database. We also make a substantial effort to annotate these signatures by parsing GEO metadata, categorized functional terms extracted by an LLM, and terms from selected Enrichr libraries. We serve these annotations alongside the gene set search results. This enables users to gain a broader perspective about the top matching gene sets. The additional metadata assists users with filtering the returned matching genes set, and identifying common themes within the top results. We also provide users with the ability to generate hypotheses for overlapping gene sets utilizing abstracts and summaries of the GEO studies, and user submitted-descriptions of their gene set. We demonstrate how this feature can uncover pathways, targets, and shared molecular mechanisms across diseases and conditions. We also show how the data within RummaGEO can be used for applications such as transcription factor and kinase enrichment analyses. Transposing the RummaGEO gene sets into a matrix that defines similarity between genes, presents the opportunity to identify gene modules and predict gene function for understudied genes. There are numerous other applications that can be enabled by reusing the RummaGEO database, for example, creating a cell type maker library for single cell identification, or developing dynamical models for cell phenotype trajectory analysis. In addition, crossing the gene sets within RummaGEO with other large sources of gene sets such as Rummagene^24^ and Enrichr^37^ we can further discover novel connections between biological processes and disease mechanisms. Overall, RummaGEO presents an unprecedented resource for the community to query, analyze, and generate hypotheses with gene expression signatures massively mined from GEO.

## Acknowledgements

This study was supported by NIH grants R01DK131525, OT2OD036435, U24CA264250, U24CA271114, and RC2DK131995.

## Data and Code Availability

The RummaGEO search engine is available from: https://rummageo.com/.

The RummaGEO data, metadata, and correlation matrices are available from the RummaGEO download page at: https://rummageo.com/download

The RummaGEO source code is available from: https://github.com/MaayanLab/rummageo.

## Competing Interests

The authors declare that they do not have any competing interests.

## Supplementary Files

**Table S1**. Gene cluster enriched functions.

**Table S2**. Key term categorizations.

## Notes

https://rummageo.com

### Competing Interest Statement

The authors have declared no competing interest.

